# Tailored Graphical Lasso for Data Integration in Gene Network Reconstruction

**DOI:** 10.1101/2020.12.29.424744

**Authors:** Camilla Lingjærde, Tonje G Lien, Ørnulf Borgan, Ingrid K Glad

## Abstract

**Background:** Identifying gene interactions is a topic of great importance in genomics, and approaches based on network models provide a powerful tool for studying these. Assuming a Gaussian graphical model, a gene association network may be estimated from multiomic data based on the non-zero entries of the inverse covariance matrix. Inferring such biological networks is challenging because of the high dimensionality of the problem, making traditional estimators unsuitable. The graphical lasso is constructed for the estimation of sparse inverse covariance matrices in Gaussian graphical models in such situations, using *L*_1_-penalization on the matrix entries. An extension of the graphical lasso is the weighted graphical lasso, in which prior biological information from other (data) sources is integrated into the model through the weights. There are however issues with this approach, as it naïvely forces the prior information into the network estimation, even if it is misleading or does not agree with the data at hand. Further, if an associated network based on other data is used as the prior, weighted graphical lasso often fails to utilize the information effectively.

**Results:** We propose a novel graphical lasso approach, the *tailored graphical lasso*, that aims to handle prior information of unknown accuracy more effectively. We provide an R package implementing the method, `tailoredGlasso`. Applying the method to both simulated and real multiomic data sets, we find that it outperforms the unweighted and weighted graphical lasso in terms of all performance measures we consider. In fact, the graphical lasso and weighted graphical lasso can be considered special cases of the tailored graphical lasso, and a parameter determined by the data measures the usefulness of the prior information. With our method, mRNA data are demonstrated to provide highly useful prior information for protein-protein interaction networks.

**Conclusions:** The method we introduce utilizes useful prior information more effectively without involving any risk of loss of accuracy should the prior information be misleading.

## Background

In the area of statistical multiomics, network models provide an increasingly popular tool for modelling complex multiomic associations and assessing pathway activity. With such models, the interactions between genes, proteins or other multiomics data can be captured and studied, and provide valuable insight into their functional relationships. This can again be used to identify pathway initiators and thus potential drug targets [1].

Networks may be constructed from data found by high-throughput gene expression profiling technologies, such as microarray or RNA-seq [2]. With the development of high-throughput multiomic technologies, large, genome-wide data sets have been made available. This enables the development of complex models integrating a variety of biological resources [3, 4]. By integrating several sources of multiomic data into a model, we can increase statistical power while providing further insight into complex biological mechanisms.

One setting where integrative network analysis has a lot of potential is when there are two types of data, e.g. measured mRNA and protein, associated with the same genes. A specific mRNA molecule is transcribed from each gene, which then can be translated into a specific protein. Thus, each gene is associated with a specific mRNA sequence and protein. With a proper model formulation, we could use information about the inferred network of one data type to improve graph inference on the other.

In this paper, we propose a novel approach to data integration in network models. The paper is organized as follows. In the remaining parts of this section, we discuss existing methodologies and the challenges we wish to address. In “Results and Discussion”, we describe our proposed methodology, and demonstrate its performance with both simulated and real multiomic data sets. We highlight our main findings in “Conclusions”, and finally give the details of our data analyses in the “Methods” section.

### Gaussian graphical network models

In a gene network model, each gene is represented by a node and an edge between two nodes represents an association between the corresponding genes. Letting each gene be associated with some measurable molecular unit (e.g. the mRNA or protein it encodes), a graph may be constructed from observed values of these *node attributes*. The attributes, each corresponding to one of *p* genes, are represented by the multivariate random vector (*X*_1_*,…, X*_*p*_)^*T*^, and a graph may be inferred from observed values of it by assuming an appropriate model.

By assuming that the vector of node attributes is multivariate Gaussian, with an unknown mean vector ***μ*** and an unknown covariance matrix **Σ**, a *partial correlation network* may be inferred by estimating the inverse covariance matrix, or *precision matrix*, **Θ** = **Σ**^*−*1^. Given the entries *θ*_*ij*_ of **Θ**, the *partial correlation* between nodes, or variables, *i* and *j* conditioned upon all others is given by

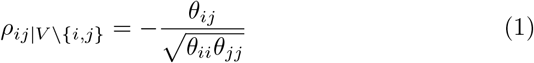

where *V* is the set of all node pairs [5]. Since correlation equal to zero is equivalent to independence for Gaussian variables, a conditional independence graph may be constructed by determining the non-zero entries of the precision matrix **Θ** and assigning edges to the corresponding node pairs. The resulting model is a *Gaussian graphical model*, with the edges representing conditional dependence.

In the Gaussian graphical model framework, the edges are assumed to be undirected and unweighted. Under these assumptions, the likelihood of the data may be derived [6]. We let ***X*** be the *n* × *p* matrix of observed data, with each row corresponding to one of *n* observations of the multivariate random vector of attributes. Letting ***S*** be the empirical covariance matrix, **Θ** then has the following profile log-likelihood:

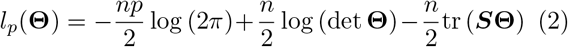

where tr denotes the trace and det the determinant. The maximum likelihood estimate for **Θ** is then the solution to the problem

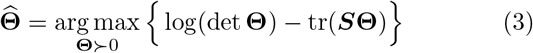

where **Θ** ≻ 0 is the requirement that **Θ** is positive definite.

In the graphical lasso, the graph is also assumed to be *sparse*. This means that it has a small edge-to-node ratio, or that the precision matrix has mostly zero elements. The sparsity of a graph can be measured by the number of edges *N*_*e*_ relative to the number of possible edges, given by 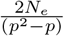.

We can not expect estimated precision matrix elements to be exactly equal to zero for real data. Further, in high-dimensional settings where the number of observations is much smaller than the number of elements to estimate, the empirical covariance matrix is not of full rank and so its inverse is not even possible to estimate directly. Thus, dimension reduction is necessary to achieve sparsity, and to estimate the precision matrix.

### The graphical lasso

*The graphical lasso* performs sparse precision matrix estimation by imposing an *L*_1_ penalty on the matrix entries [6]. It is constructed to solve a penalized version of the log-likelihood problem (3),

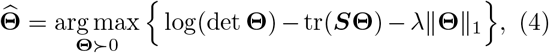

where ∥ · ∥_1_ is the *L*_1_ norm and *λ* a penalty parameter that must be tuned [7]. Due to the *L*_1_ penalty, the method sets many elements of **Θ** to zero. *λ* controls the sparsity, and a larger value of it results in fewer included edges. Utilizing the fact that solving problem (4) is equivalent to iteratively solving and updating a lasso least-squares problem [7, 8], the graphical lasso algorithm is both exact and computationally efficient [6].

The method is implemented in the R packages `glasso` [9] and `huge` [10], with the latter providing several routines for penalty parameter selection.

### Penalty parameter selection

There are several penalty parameter selection methods available, and we have used two of the more common ones in our analyses.

#### StARS

Stability Approach to Regularization Selection (StARS) is a selection method based on model stability [11]. The method starts with a large penalty *λ* corresponding to an empty graph, i.e. a graph with no edges, and decreases it stepwise. For each value of *λ* many random subsamples are drawn from the data, and the graphical lasso is used to fit a graph for each sample. As a measure of the instability of each edge under the sub-sampling, the average number of times any two graphs disagree on the edge is found. By averaging this instability measure for all edges, the total instability is found.

Given a cut point *β* for the instability we are willing to accept, the corresponding *λ* is selected as the optimal penalty parameter. *β* may be interpreted as the fraction of edges we are willing to accept as possibly wrong, and it is normally set to 0.05. This way, StARS aims to choose the least amount of regularization that makes graphs sparse as well as reproducible under random sampling. It should be noted that since the method constructs a graphical lasso graph for every subsample, it is computationally costly for large graphs.

#### The extended BIC

The extended BIC (eBIC) is a modified version of the Bayesian Information Criterion constructed for selection in high-dimensional graph settings [12]. For a given edge set *E*, it is given by

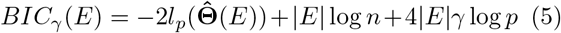

where |*E*| is the number of edges in the edge set *E* and where γ ∈ [0, 1]. If γ = 0 we get the ordinary BIC, while positive values give stronger penalization of large graphs. The parameter γ can only be tuned by experience, and should be large enough to get a low false discovery rate, but small enough to get a satisfactorily high positive discovery rate [12].

Like for the ordinary BIC, the penalty corresponding to the model that minimizes the eBIC is chosen. This selection criterion is computationally efficient, and has been shown to outperform both the ordinary BIC and cross validation when the sizes of *p* and |*E*| are comparable to *n*. This holds for any choice of γ ∈ (0, 1] [12]. In our applications, we are comparing relatively small graphs where the extra penalization due to high-dimensionality is not as needed. We are comparing graphs of similar sparsity, which also means that the extra penalization of edges is not that important. In our simulations we are therefore simply choosing γ = 0 so that we get the ordinary BIC, but for generality we propose to use the eBIC criterion in our method.

### The weighted graphical lasso

The weighted graphical lasso is an extension of the graphical lasso which allows the incorporation of additional information through a *p* × *p* weight matrix ***W*** with entries in [0, 1] [13]. Theoretically, this can be justified by the Bayesian interpretation of the graphical lasso, and ***W*** can be regarded as a *prior matrix* representing prior information about the existence of edges [13]. Using the *penalty matrix **P*** = 1 - ***W***, the estimated precision matrix must now satisfy

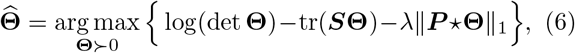

where ⋆ denotes component-wise matrix multiplication. (6) can be regarded as the problem (4) with prior information incorporated into the expression. It is clear that while an edge with prior weight equal to zero is not penalized at all, no edge is penalized by a factor larger than *λ*. Thus, edges with the minimal penalty are almost guaranteed to be included while edges with the maximum penalty are not necessarily guaranteed to be excluded, unless *λ* is infinitely large.

Similarly to the original problem, optimization of the problem (6) may be done using the graphical lasso algorithm. The modification can easily be incorporated into the algorithm by replacing *λ* by individual penalties 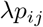, *j* ≠ *i*, where *p*_*ij*_ is the *ij*^th^ element in ***P***.

If the prior information is informative, the weighted graphical lasso is found to outperform the graphical lasso in simulations [13]. However, there are several potential issues with the weighted graphical lasso. Firstly, it does not take the possibility of the prior information being partly or totally misleading into account. If an excess amount of prior information is incorrect, it can be harmful to the final estimates [13]. For example, edges corresponding to zero entries in ***P*** are not penalized and therefore almost guaranteed to be included in the final model, even if this is not supported by the data.

Secondly, the weighted graphical lasso might not be able to differentiate enough between weights. The range and distribution of the weights, and thus the penalties, is not necessarily purposeful, often leading to limited effect of otherwise valuable prior information. In such a case it could be sensible to use a non-linear transformation of the weights. This is especially important if the prior weights do not have a linear interpretation where having twice the weight indicates having twice the confidence in an edge, which could be the case if the prior weight matrix is found from the estimated precision matrix of another, related data set [13]. On the other hand, it could be that the prior information is partly informative in that it is only useful to include it to a limited degree. It is altogether clear that the prior weights should be given more consideration, and not just used naïvely in the weighted graphical lasso procedure as in [13].

## Results and discussion

### The tailored graphical lasso

To deal with the shortcomings of the weighted graphical lasso, we propose a novel approach that aims to handle prior information of unknown accuracy and to utilize it more effectively. The idea is to use a nonlinear transformation *g*_*k*_(*w*) of the weights, where the behavior of the function *g*_*k*_(*w*) is controlled by a parameter *k* with parameter space Ω. We may then choose the best transformation in the function space 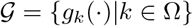 by considering the precision matrix estimates we obtain for various values of *k* in a suitable partition of Ω. The optimal value *k*_opt_ is chosen using some selection criterion. Ideally both the identity function *g*_*k*_(*w*) = *w* and the zero function *g*_*k*_(*w*) = 0 should be contained in 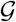 so that both the ordinary weighted graphical lasso and the ordinary graphical lasso is considered. This way, we attempt to avoid transformations that may result in worse estimates than these two standard methods.

Inspired by the ideas of [4], we propose a logistic function for the weight transformation. Figure 1 shows the logistic function 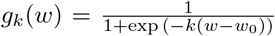, where *w*_0_ is the sigmoid midpoint and *k* the steepness parameter. Evidently, for *k* = 0 the function maps all weights to the same value, and we just get the ordinary unweighted graphical lasso. As *k* grows, the sigmoid function becomes more step-like. For *k* = 40 it is very near being a step function and for *k* = 200 it essentially is one. The transformed weights will always be mapped into [0, 1], as required for the weights in the weighted graphical lasso. The logistic function never becomes exactly the identity function *g*_*k*_(*w*) = *w* that would map all weights to themselves corresponding to the ordinary weighted graphical lasso. As seen from Figure 1, it does however approximate the identity function well for *k* = 4.

**Figure 1.**
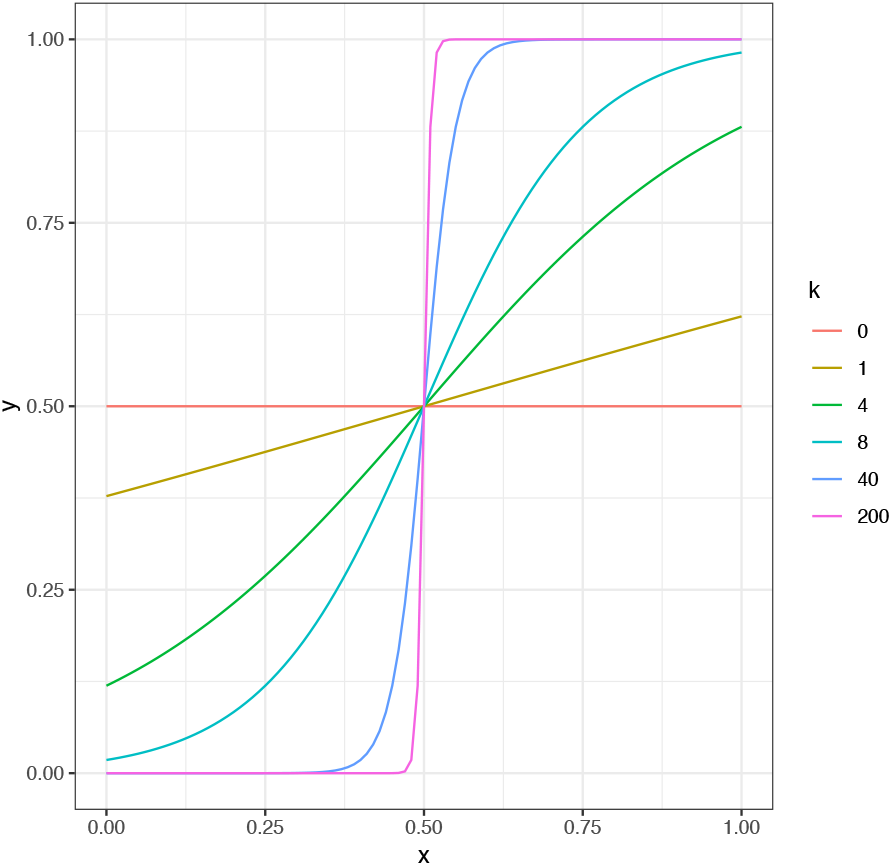
The logistic function for *w*_0_ = 0.5 and various values of the steepness parameter *k*.

Thus, by appropriate data-driven tuning of the parameter *k* we may interpret a small selected *k* as an exclusion of the prior information and a large one as an inclusion and enhancement of it, giving the edges weights close to 0 or 1 depending on whether their prior weights are below or above the threshold *w*_0_. This interpretability is convenient, as the optimal value of *k* found by some selection criterion then tells us how useful the prior weights are and whether increasing their differences even more improves the model.

Using the logistic weight transformation, we propose a method we call *the tailored graphical lasso*. In the algorithm, we begin by selecting the total amount of penalization. We do this with StARS, using the unweighted graphical lasso to find a common penalty parameter *λ*. For each *k*, we find the matrix ***W***_*k*_ of weights transformed with the logistic function with this steepness parameter, which gives us the penalty matrix ***P***_*k*_ = 1 − ***W***_*k*_. We then choose the penalty parameter *λ*_*k*_ in the tailored graphical lasso that preserves the amount of penalization selected for the unweighted graph, i.e. so that *λ*_*k*_∥***P***_*k*_∥_1_ = *λp*^2^. We use StARS as this is a stable selection criterion for graph sparsity selection [11]. By preserving the amount of penalty, we achieve similar sparsity for each *k* without having to repeatedly perform StARS, which is computationally exhaustive.

As a criterion for choosing *k*, we use the eBIC as it is computationally efficient. In the criterion, we choose a value of γ ∈ [0, 1] that reflects how concerned one is with false discoveries. The larger it is, the more we penalize larger graphs. As for the sigmoid midpoint *w*_0_, we let it be equal to the lower *β*-quantile of the non-zero prior weights, where *β* is the variability threshold used in the StARS tuning of the ordinary graphical lasso graph. This is usually set to a default value of 0.05. This way, we avoid having to tune a second parameter. We justify our choice of *w*_0_ by the fact that *β* is the upper limit set in the StARS selection for the estimated probability of an inferred edge being wrong. This means that we can expect up to a fraction *β* of the inferred edges of the final graph as tuned by StARS to be incorrect.

The algorithm is shown in Table 1. After using the method, the common penalty parameter 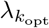 might be adjusted slightly to achieve the exact sparsity found to be optimal in step 1. The maximum value *k*_max_ must also be chosen, and as visible from Figure 1 it is sufficient to choose it to be 80, as the logistic function basically is a step function at that point.

**Table 1.**
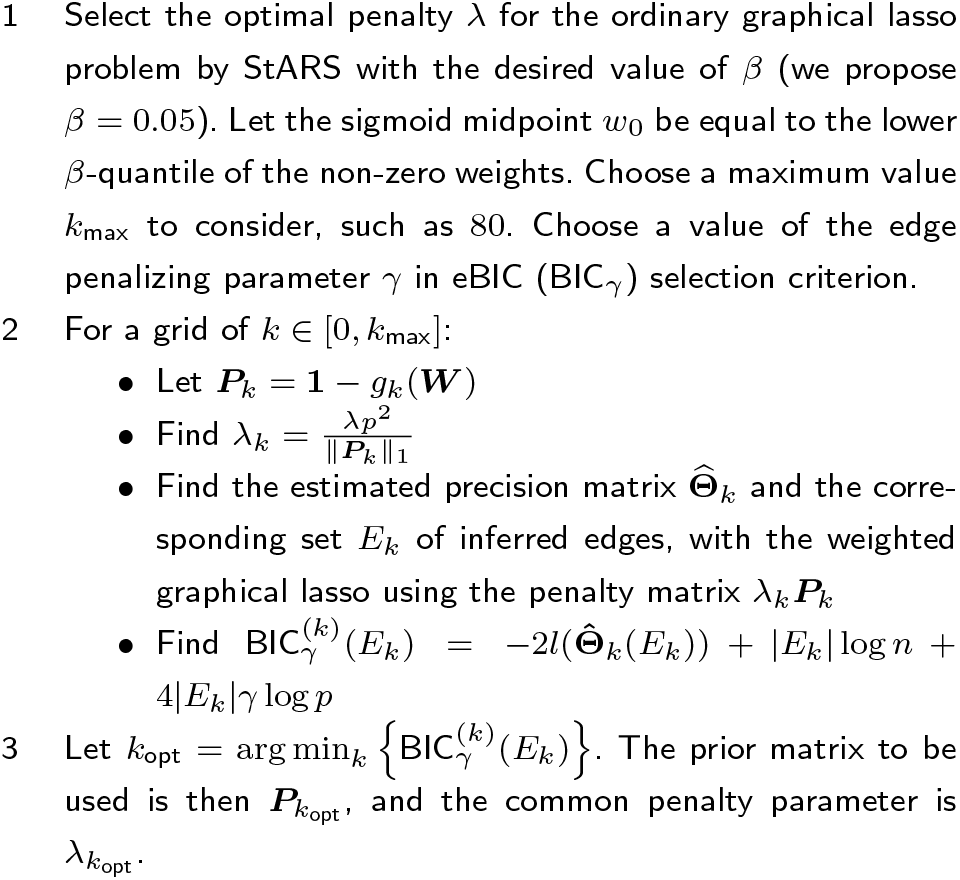
The tailored graphical lasso algorithm.

We have implemented the tailored graphical lasso in the R package `tailoredGlasso` (https://github.com/Camiling/tailoredGlasso).

### Simulated data

To evaluate the performance of the tailored graphical lasso, we have done comprehensive simulation studies in R [14]. The details are given in Methods. We have used our R package `tailoredGlasso` to perform the tailored graphical lasso, and the code for the data analysis is available on Github (https://github.com/Camiling/tailoredGlassoAnalysis).

To make our simulations as relevant to our multiomic application of interest as possible, we have generated data with the *scale-free property*, which is a known trait in multiomic data [5]. We have simulated various sets of data with the same “true” underlying graph structure, with a sparsity of 0.02 and *p* = 100 nodes. We let the sizes of the partial correlations vary between data sets. The data sets are generated from the corresponding multivariate Gaussian distributions. Letting there be *n* = 80 observations in each data set, we have a high-dimensional problem with (*p*^2^ − *p*)*/*2 = 4950 potential edges. The graph structure is shown in Figure 2 with nodes colored according to their *degree*, the number of adjacent edges, where darker color indicates higher degree.

**Figure 2.**
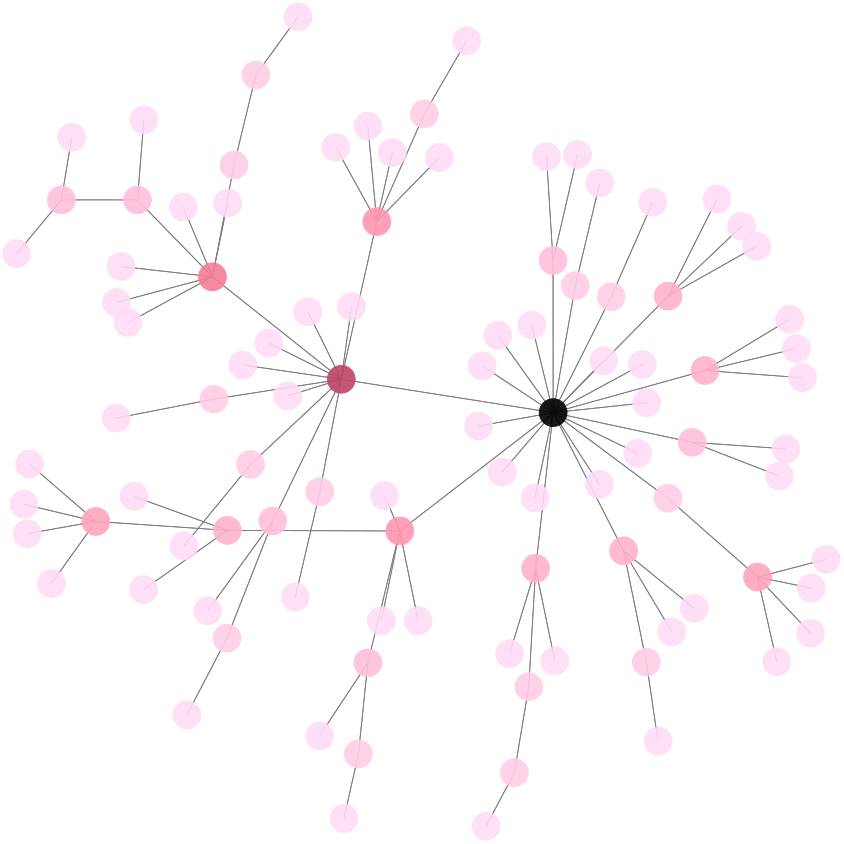
Graph structure of simulated data.

For each data set, we have also generated various prior weight matrices in order to investigate the performance of the method in different settings. Specifically, we have created *prior precision matrices* of various similarities to the precision matrices of interest, generated *prior data* from the corresponding multi-variate Gaussian distribution and used the ordinary graphical lasso to estimate the prior precision matrix. A prior weight matrix is then created by taking the absolute values of the corresponding partial correlation estimates using the formula (1). The prior matrices are constructed this way to mimic real applications where we have two related data sets, both with unknown precision matrices, and we want to use information from one to improve the precision matrix estimate of the other. The precision matrix of the data used as the prior then needs to be estimated, and a prior weight matrix is constructed from it [13]. These are the prior matrices we use in the weighted graphical lasso and the tailored graphical lasso.

In our simulations, we have considered 7 different combinations of partial correlations in the network of interest and the prior, and the fraction of edges that the network of interest and the prior network disagree on. This way, we get several prior precision matrices of various accuracy and various strength of the partial correlations. For each of these cases, after the data has been generated, we have used both the unweighted and the weighted graphical lasso in addition to our proposed tailored graphical lasso to estimate the precision matrix and reconstruct the underlying graph structure. We have for each modified prior simulated *N* = 100 corresponding data sets, and averaged the results when the above methods were applied.

The estimated graphs are assessed by the *precision*, which is the fraction of the inferred edges that are actually present in the true graph, and *recall*, which is the fraction of the edges in the true graph that are present in the inferred one. While both measures are important in understanding how well a graph reconstruction method has estimated a graph, we put more emphasis on the precision as we are more concerned about false positives that false negatives. In a multiomic setting, we would not want to identify many inaccurate interactions, but rather identify fewer but more trustworthy ones. As opposed to the precision, the recall will necessarily grow as more edges are included, and does not tell us how accurate the estimated edges that are present are.

Table 2 shows the averaged results for the different cases, where we have abbreviated the graphical lasso and weighted graphical lasso as glasso and wglasso, respectively. As expected, we see that the selected *k*, i.e. *k*_opt_, increases with the accuracy of the prior weights.

**Table 2.** Performance of different graph reconstruction methods in simulations. The performance of the different graph reconstruction methods in simulations. The edge disagreement between the graph of interest and its prior, as well as the size of the partial correlations in them, is shown as well. The results are averaged over *N* = 100 simulations. The best values of the different performance measures are marked in bold, and *k*_opt_ is the mean value of the *k* chosen by the eBIC selection criterion in the tailored graphical lasso (TailoredGlasso). The graphical lasso and weighted graphical lasso are abbreviated as Glasso and Wglasso, respectively.

| Case | Edge disagreement % | Partial cor | Prior partial cor | Method | $k_{\text{opt}}$ | Sparsity | Precision | Recall |
| --- | --- | --- | --- | --- | --- | --- | --- | --- |
| 1 | 0 | 0.2 | 0.2 | Glasso | - | 0.035 | 0.283 | 0.493 |
|  |  |  |  | Wglasso | - | 0.032 | 0.312 | 0.503 |
|  |  |  |  | TailoredGlasso | 49.64 | 0.031 | **0.389** | **0.606** |
| 2 | 0 | 0.2 | 0.1 | Glasso | - | 0.035 | 0.285 | **0.499** |
|  |  |  |  | Wglasso | - | 0.034 | 0.293 | 0.493 |
|  |  |  |  | TailoredGlasso | 13.39 | 0.034 | **0.295** | 0.496 |
| 3 | 0 | 0.1 | 0.2 | Glasso | - | 0.022 | 0.079 | 0.085 |
|  |  |  |  | Wglasso | - | 0.021 | 0.096 | 0.099 |
|  |  |  |  | TailoredGlasso | 3.34 | 0.021 | **0.100** | **0.103** |
| 4 | 0 | 0.1 | 0.1 | Glasso | - | 0.022 | 0.079 | **0.085** |
|  |  |  |  | Wglasso | - | 0.02 | 0.082 | 0.083 |
|  |  |  |  | TailoredGlasso | 5.63 | 0.02 | **0.083** | 0.084 |
| 5 | 10 | 0.2 | 0.2 | Glasso | - | 0.035 | 0.283 | 0.493 |
|  |  |  |  | Wglasso | - | 0.033 | 0.305 | 0.493 |
|  |  |  |  | TailoredGlasso | 41.2 | 0.032 | **0.335** | **0.532** |
| 6 | 20 | 0.2 | 0.2 | Glasso | - | 0.035 | 0.283 | **0.493** |
|  |  |  |  | Wglasso | - | 0.034 | **0.291** | 0.491 |
|  |  |  |  | TailoredGlasso | 4.31 | 0.034 | **0.291** | **0.493** |
| 7 | 100 | 0.2 | 0.2 | Glasso | - | 0.035 | 0.283 | **0.493** |
|  |  |  |  | Wglasso | - | 0.034 | 0.289 | 0.485 |
|  |  |  |  | TailoredGlasso | 2.94 | 0.034 | **0.290** | 0.485 |

For the most accurate prior network (case 1), *k* is selected to be as large as 49.64, and we see that the precision of the tailored graphical lasso is 37% higher than for the graphical lasso and 25% higher than for the weighted graphical lasso. In this case the recall is also 23% higher than for the graphical lasso and 20% higher than for the weighted graphical lasso. We note that the ordinary weighted graphical lasso does not perform much better than the unweighted graphical lasso. The reason is that the absolute values of partial correlations for the prior data are between 0 and 0.2, so the inclusion of these as prior weights in the ordinary weighted graphical lasso does not affect the resulting estimates too much. But by using a logistic transformation, we are able to enhance the difference of the prior weights, and this results in a much improved performance of the tailored graphical lasso

On the other hand, for completely misleading prior weights (case 7), *k* is small for the tailored graphical lasso and we get results similar to the ordinary graphical lasso. The fact that the optimal *k* is not chosen to be exactly zero can be due to randomness, where the weak inclusion of the prior information actually improves the inference.

In the other cases, the optimal *k* is larger and the tailored graphical lasso either outperforms or performs as well as the two other methods in terms of the precision.

### Multiomic data

For illustratory and investigatory purposes, we have applied the tailored graphical lasso to real multiomic data sets. Comparing the results to those from the ordinary weighted graphical lasso, we want to see if the tailored lasso better fits the data and can identify more multiomic interactions for which there is evidence in the literature.

We have used two data sets, the first one containing *n* = 743 breast cancer tumor samples from the well known TCGA BRCA database [15]. We considered gene expression measured by RNA-seq for *p* = 165 genes, as well as their associated protein measured by Reverse Phase Protein Array (RPPA). We have only considered the genes known to encode the proteins present in the RPPA data panel. The other data set is called Oslo 2, and consists of *n* = 280 breast cancer samples collected from hospitals in Oslo [16]. We have considered gene expression measured by microarray for *p* = 100 genes, as well as their associated protein measured by RPPA.

While the research done on genomic networks using mRNA measures already is quite substantial, the field of protein-protein interaction networks is newer. It is therefore interesting to see if the large knowledge learned by gene expression analysis can be used in the construction of a protein network. Thus, we will for each of the two data sets infer protein-protein interaction networks from the RPPA data, treating the mRNA data as prior information.

The prior weights are constructed by using the graphical lasso to estimate the precision matrices of the mRNA data, and letting the weights be the absolute values of the corresponding partial correlations. The partial correlation between two genes gives us the correlation between them when conditioning upon all other gene interactions, and this approach results in less penalization of edges between proteins whose associated genes are found to have a strong relationship in the corresponding mRNA network. The resulting partial correlation weights are in the range [0, 0.2], just as for our simulated data. It should be noted that for genomic data, partial correlations will indeed tend to lie in this range and so a partial correlation of 0.2 can be considered large (see for example [17–19]).

The distributions of the prior weights of the simulated and real multiomic data are also very similar, which also indicates that our simulations were close to the application of interest (see Additional file 1).

After the precision matrices of the RPPA data are estimated using the different methods, we assess how well they fit the data by computing the corresponding multivariate Gaussian log likelihoods (2). Since this log likelihood will depend on the sparsity of the estimated graph, and generally increases with the number of included parameters (edges), we have forced the graphs estimated by the different methods to have the same sparsity as was found optimal by StARS on the unweighted graph. This way, direct comparison is possible.

Table 3 shows the results for the TCGA data. As we see, the log likelihood of the estimated precision matrix is indeed largest for the tailored graphical lasso estimate. The results for the Oslo 2 data set is shown in Table 4, and also here the tailored graphical lasso estimate has the largest log likelihood. For both data sets, the increase in the log likelihood when comparing the tailored graphical lasso to the weighted graphical lasso is much larger than when comparing the weighted graphical lasso to the ordinary graphical lasso.

**Table 3.** Comparison of results for the TCGA data set, with the highest log-likelihood value in bold.

| Method | $k_{\text{opt}}$ | Sparsity | $l_p(\hat{\Theta})$ |
| --- | --- | --- | --- |
| Glasso | - | 0.034 | -162226.8 |
| Wglasso | - | 0.034 | -161928.4 |
| TailoredGlasso | 127.00 | 0.034 | **-160697.1** |

**Table 4.** Comparison of results for the Oslo 2 data set, with the highest log-likelihood value in bold.

| Method | $k_{\text{opt}}$ | Sparsity | $l_p(\hat{\Theta})$ |
| --- | --- | --- | --- |
| Glasso | - | 0.026 | -38489.65 |
| Wglasso | - | 0.026 | -38419.44 |
| TailoredGlasso | 149.0 | 0.026 | **-38132.04** |

The optimal *k* selected by the tailored graphical lasso is very large in both cases, meaning the method finds that the information from the mRNA networks improves the inference on the protein networks. For such large values of *k*, prior weights are mapped to values close to either 0 or 1, depending on whether they are above or below the sigmoid midpoint *w*_0_. This is an interesting result, as it means that the mRNA networks is found to provide very useful information about the protein networks.

To investigate whether the edges in the graphs found by the tailored graphical lasso have more evidence in the literature than the ones found by the ordinary weighted graphical lasso, we have also performed data mining using the STRING database of known and predicted protein-protein interactions [20]. Focusing on the edges the tailored graphical lasso and the weighted graphical lasso disagree on, we have found the fraction of edges present only in the tailored graphical lasso graph that have proof in the STRING database, and compared it to the fraction for the ones present only in the weighted graphical lasso graph.

The two methods disagreed on 296 edges in the TCGA data, and 103 edges in the Oslo 2 data, which means that they disagree on the majority of the edges. Table 5 shows the percentage of the edges unique to the different graphs that have evidence in the STRING database. As we see, for both data sets there is far more evidence for the edges unique to the tailored graphical lasso graphs than those unique to the weighted graphical lasso graphs.

**Table 5.** Comparison of evidence for edges unique to each graph. Comparison of evidence for edges unique to each graph, using the STRING database. The highest percentage of edges with evidence is in bold.

| Data set | Edge evidence % |  |
| --- | --- | --- |
|  | Wglasso | TailoredGlasso |
| TCGA | 18.0% | **32.2%** |
| Oslo 2 | 28.2% | **43.7%** |

The resulting optimal graphs are shown in Additional files 2 and 3 for TCGA and Oslo 2 data sets respectively. It is reassuring that the essential breast cancer genes are central (hubs) in the resulting networks, such as the tumor suppressor TP53 and the estrogen receptor ESR1, as we would expect [21, 22]. To investigate the nature of the identified interactions, we have taken a closer look at the type of evidence they have in STRING. The STRING database gives each interaction it identifies a score for each type of evidence it considers. The types of evidence include experimentally determined interactions, coexpression determined by large-scale analyses, annotated pathways found from data bases and literature mining of co-occurrence. The scores lie in [0, 1] and are indicators of confidence, i.e. how likely STRING judges an interaction to be true, given the available evidence [23]. A score equal to 1 indicates the highest possible confidence while a score of 0.5 would indicate that roughly every second interaction might be erroneous. In STRING, a score ≥ 0.4 is considered “medium confidence”, and this threshold is proposed for determining relevant interactions [20].

For both the Oslo 2 and the TCGA data sets, we have found the total number of interactions with edge score ≥ 0.4 for each type of evidence in STRING. The results are shown in Additional file 8. We find that for the different types of evidence, the number of edges with at least medium confidence is larger for the tailored graphical lasso graphs than for the classical weighted graphical lasso graph. This is expected since the tailored graphical lasso identified more edges with evidence in STRING overall. In particular, the tailored graphical lasso graphs had far more edges with evidence from coexpression analyses and literature mining. As for the other evidence type such as experimentally determined interactions and annotated pathways, the tailored graphical lasso graphs also had more edges with evidence in STRING than the weighted graphical lasso, but the difference was not as large.

Lists of all gene pairs with interactions in the tailored graphical lasso graphs of the data sets are given in Additional files 4 and 5. Lists of the gene interactions that the tailored graphical lasso was able to find, but not the weighted graphical lasso, are given in Additional files 6 and 7.

## Conclusions

In this paper, we introduce the tailored graphical lasso as an extension to the weighted graphical lasso for graph reconstruction. The objective is to get better utilisation of the available prior information, while ensuring that the introduction of prior information may not decrease the accuracy of the resulting inferred graph. The method is implemented in the R package `tailoredGlasso`.

The method is developed with multiomic applications in mind, and to illustrate the performance of the method in such settings we have simulated data similar to this application. We have considered different scenarios in which the strength of the partial correlations varies both in the network of interest and the prior network, and in which the edge agreement between these networks varies. We found that if the prior information is completely useless and its inclusion in the weighted graphical lasso only results in a less accurate graph estimate, the tailored graphical lasso will give results similar to the ordinary graphical lasso and weighted graphical lasso.

On the other hand, if the prior information is informative, the tailored graphical lasso will outperform the graphical lasso and perform either as well as or better than the weighted graphical lasso. For less useful prior information the two methods will perform very similarly, and as the usefulness of the prior information increases the tailored graphical lasso will have better results as it utilizes the prior information more effectively.

The method also has a nice interpretability through the estimated value of *k*, giving us a “usefulness score” for the prior information, where *k* close to zero indicates that the prior information does not provide any useful information while larger *k* indicates that it does. *k* > 0 means that inclusion of the prior information to some degree is found to be useful, and *k >* 4 means that enhancing the differences in the prior weights is found to be beneficial. Both for the TCGA and the Oslo 2 data set, *k* was found to be very large (127 and 149, respectively), meaning that the mRNA data was indeed found to be an informative prior for the less studied protein data.

We have applied the method to multiomic data from two studies on breast cancer, TCGA BRCA and Oslo 2, showing that mRNA data can be useful as prior information for protein-protein interaction networks. The estimated precision matrices found by the tailored graphical lasso had higher log likelihoods than the ones found by the ordinary weighted and unweighted graphical lasso. Further, through data mining with the STRING database we found that there is indeed more evidence for the graph structures found by the tailored graphical lasso than the two other methods, in particular evidence in the form of co-occurrence in literature and co-expression in large-scale analyses.

Altogether, we have seen that the tailored graphical lasso performs either as well as the ordinary weighted and unweighted graphical lasso, or better, depending on the usefulness of the prior information. This means that there is much to gain yet little to lose from using the tailored graphical lasso, as it allows us to use priors of unknown accuracy without taking risks.

## Methods

### Data simulations

#### Generating Gaussian graphs and data

In our multiomic application, the weighted graphical lasso is performed by first using the graphical lasso on the mRNA data sets and RPPA data sets separately, and then using the estimated mRNA network as a prior network for the RPPA one. This is done by letting the prior weights be the absolute values of the resulting estimated partial correlations of the mRNA networks. Those estimates are found from the estimated precision matrix of the mRNA data, using formula (1). In our simulations we aim to mimic this setting, and so we will for each simulated set of data create a similar, but not identical, set of prior data.

We have done our simulations in R using the package `huge`. In particular, we have used the function `huge.generator()`, which allows us to generate data with the *scale-free property* as is a known trait in multiomic data [5]. We let all networks consist of *p* = 100 vertices, and let there be *n* = 80 observations of the corresponding multivariate data. The final graph has *p* edges and thus a *sparsity* of approximately 0.02. As for the values of the non-zero partial correlations, we let them all be equal to either 0.1 or 0.2.

Once a precision matrix **Θ** is constructed, huge generates the multivariate Gaussian data of the vertices from the distribution with covariance matrix **Σ** = **Θ**^*−*1^ and expectation vector 0.

#### Generating prior data

For a set of simulated data, we have modified its precision matrix and generated data from the resulting distribution. This way, we get sets of prior data from distributions of various similarity.

Specifically, for an initial precision matrix we have created, we have permuted a certain fraction of the edges by randomly redirecting all the edges of some nodes to others. We then consider each of those permuted precision matrices with all partial correlations being either 0.1 or 0.2, resulting in precision matrices of various accuracy and various strength of the partial correlations.

For each prior precision matrix **Θ**_prior_, a prior data set ***X***_prior_ is generated from the resulting multivariate Gaussian distribution. The graphical lasso is then used on this data to obtain a prior precision matrix estimate 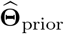, which is used to construct a prior matrix by finding the absolute values of the corresponding partial correlation estimates.

#### Data analysis

For each combination of precision matrix and prior data, a precision matrix estimate 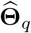 for the data ***X*** of interest is found for each of the weight-based procedures of the tailored graphical lasso and the weighted graphical lasso. An ordinary unweighted graphical lasso estimate 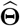 of the data is also found for comparison. The whole data simulation procedure is shown in Figure 3.

**Figure 3.**
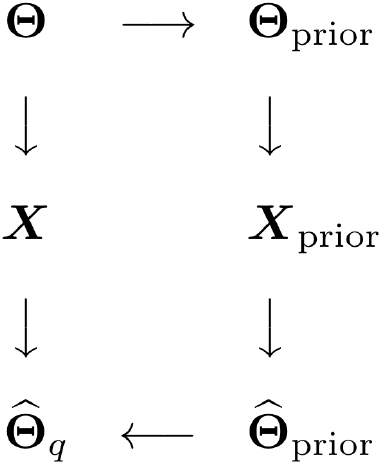
Illustration of data generation and analysis procedure in simulations. Illustration of data generation and analysis procedure in simulations. For each precision matrix **Θ** we consider, we modify it to create a prior precision matrix **Θ**_prior_. A prior data set ***X***_prior_ is generated from the resulting multivariate Gaussian distribution. The graphical lasso is then used on this data to obtain a prior precision matrix estimate 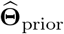, and the absolute values of the corresponding partial correlation estimates are used as weights in the tailored and weighted graphical lasso to get a precision matrix estimate 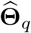 for the data ***X*** of interest.

The ordinary graphical lasso was performed using the R function `huge`, while the weighted graphical lasso was performed using the `glasso` function in R as it allows for the use of a prior matrix in the penalty. We have tuned the weighted and unweighted graphical lasso graphs by StARS with instability threshold *β* = 0.05, and according to our proposed algorithm we choose the sigmoid midpoint *w*_0_ as the lower 0.05-quantile of the non-zero prior weights in the tailored graphical lasso. For simplicity, in the tailored graphical lasso we have used the eBIC with parameter *γ* = 0 as our selection criterion. This is justifiable as we are comparing graphs of very similar sparsity, and so the extra penalization of the number of edges in the graphs is not necessary. The results are averaged over *N* = 100 simulations for all cases. The quantities we report are the precision and recall, as well as the sparsity of the estimates and the chosen *k* in the tailored graphical lasso.

### Multiomic data

Below we give a description of the preprossessing of the two data sets we have used, before describing the analysis we have done.

#### TCGA BRCA

The breast cancer tumor data from the TCGA BRCA database was downloaded from the UCSC Xena Browser [24]. We have used *n* = 743 breast cancer tumor samples, considering gene expression measured by RNA-seq for *p* = 165 genes as well as their associated protein measured by Reverse Phase Protein Array (RPPA). The RPPA data is log2(*x*) transformed and median centered, while the RNA-seq data has been normalized by upper quartile FPKM and log2(*x* + 1) transformed [15]. The full data set is larger than the subset we have used, but to make the data applicable to our methods a reduction was necessary. First, we have disregarded “control” samples in the data set, since our interest is in the samples linked to breast cancer. We have also disregarded phosphorylated proteins in the RPPA data set, as they are only a subgroup of the full measured proteins.

Some proteins have been measured by antibodies taken from different animals, meaning some protein measures in the data set are actually for the same protein. They are complimentary in their missingness in the sense that samples only have a measurement for one of them, and to get the complete protein profiles we therefore need to merge these values. This is done by taking their mean values after missing ones are removed. Further, there are 137 samples with many missing RPPA measurements that we discard. There are also 4 proteins whose values are missing in almost all samples, and these have been removed from the data set as well.

The full RNA-seq data set includes gene expression measurements for 60, 483 genes, but we have only considered the genes known to encode the proteins present in the RPPA data set. Some samples have been split into several vials, and for those we have chosen to use vial A. Finally, the gene XBP1 was found to have all RNA-seq values equal to zero, and so we have removed this gene and its corresponding protein from the data set. This is necessary as the graphical lasso does not allow zero-variance variables.

To get a complete one-to-one correspondence between the two data types we chose to only look at the samples that were present in both data sets. Before the final analysis, the variables in the data sets were scaled and centered across samples to ensure scale-invariant penalization in the graphical lasso-based methods.

Finally, the mapping between the proteins and the associated genes that encode them was found mainly from the UCSC Xena Browser, however, not all proteins aliases were represented there. Therefore, mappings from the Stanford-Cancer Genome Atlas database [25] and the GeneCards database [26] were also used.

#### Oslo 2

The Oslo 2 data we have used includes *n* = 280 breast cancer samples collected from hospitals in Oslo [16]. The gene expression for *p* = 100 genes has been quantified by measuring mRNA with microarray-technology, and their associated proteins have been measured by RPPA technology. The data set containing the mRNA measurements was downloaded from the GEOquery database using the Bioconductor package in R [27], while the protein measurements are downloaded from the eprint version of the paper “Integrated analysis reveals microRNA networks coordinately expressed with key proteins in breast cancer” [16]. The microarrays are by default log2(*x*) transformed, quantile normalized and hospital adjusted by subtracting from each microarray probe value the mean probe value among samples from the same hospital. This transformation and normalization is commonly done on multiomic data to get approximately normal data, as is required for the graphical lasso to work properly.

Like for the TCGA BRCA data set, some modifications of the Oslo 2 data set were necessary. First, we have only included genes in the mRNA data set that encode proteins present in the RPPA data set. To avoid violation of the independence assumption, in the case of a patient having several tumor samples from different pathological areas we have only considered the first one. Further, the measurements for two of the antibodies in the RPPA data were removed as they are known to bind to several proteins.

We have also merged some mRNA measures in the cases where there are several probes known to bind to the same gene. These are highly correlated, and so merging them is plausible. This is done by taking the mean of the scaled and centered measurements. The scaling ensures that the result is independent of the scale of the different antibodies.

Finally, the final data sets are scaled and centered, so that each gene or protein expression measurement has mean 0 and standard deviation 1 across samples.

#### Analysis

For both the TCGA BRCA and the Oslo 2 data set, we have inferred protein-protein interaction networks using both the weighted graphical lasso and the tailored graphical lasso. For each data set, the mRNA data is treated as prior information, letting the prior weights be the absolute values of its estimated partial correlations. The estimated partial correlations for the mRNA data are found from the graphical lasso estimate for the precision matrix using the formula (1).

The graphical lasso on the mRNA data was performed using the huge package in R, using StARS with instability threshold *β* = 0.05 to select the penalty parameter. According to our tailored graphical lasso algorithm, we then choose the sigmoid midpoint *w*_0_ as the lower 0.05-quantile of the non-zero prior weights in the tailored graphical lasso. We have used the eBIC with parameter *γ* = 0.6 as our selection criterion in the tailored graphical lasso. We chose this larger value to reflect that we are more concerned about false positives than false negatives, as we will interpret the genomic implications of the results. However, as previously discussed we are comparing graphs of very similar sparsity, and so the choice of extra penalization of the number of edges in the graphs will not make a big impact.

As for the parameter *k*, we find it sufficient to consider a grid of values in [0, 150] as *k* necessarily is non negative, and since the logistic function essentially has the same step-function shape for all *k* larger than 100 as shown in Figure 1. The weighted graphical lasso was performed using the `glasso` function in R, as it allows for the use of a prior matrix in the penalty.

After the precision matrices of the protein data of each data set is estimated using the different methods, we assess how well they fit the data by computing the corresponding multivariate Gaussian log likelihoods (2). Since this log likelihood will depend on the sparsity of the estimated graph, and generally increases with the number of included parameters (edges), we have forced the graphs estimated by the different methods to the sparsity found optimal by StARS on the unweighted graph. This way, direct comparison is possible.

#### The STRING database

While the fact that the tailored graphical lasso estimates have a higher log likelihood value than the weighted graphical lasso ones implies that the resulting graph explains the data better, it might be interesting to investigate the results even further. One possible way to check whether the new edges are more plausible is to check whether there are other sources supporting the existence of the gene relationships the edges represent. For this purpose, we have used the STRING database of known and predicted protein-protein interactions [20]. As described on the STRING website, the interactions include direct (physical) and indirect (functional) associations and are derived from five main sources, namely genomic context predictions, high-throughput lab experiments, co-expression, automated text-mining and previous knowledge from other databases.

The way we check the database for evidence of the existence of a set of edges is to feed a list of all genes involved to the STRING search engine. STRING then provides a graph where an edge between two nodes means that the database has evidence for the existence of an interaction between the two genes the nodes represent. A list of the edges in the resulting graph may then be downloaded as a.tsv file, and we can check how many of the edges in our original edge set that are present in this list. Focusing on the edges the tailored graphical lasso and the weighted graphical lasso disagree on, we may then find the fraction of edges present only in the tailored graphical lasso graph that have proof in the STRING database, and compare it to the fraction for the ones present only in the weighted graphical lasso graph. If the latter fraction is lower, the edges found by the tailored graphical lasso method have more support in the database.

## Abbreviations

mRNA: Messenger Ribonucleic Acid
RPPA: Reverse Phase Protein Array
eBIC: Extended Bayesian Information Criterion
StARS: Stability Approach to Regularisation Criterion

## Acknowledgements

Not applicable.

## 1 Authors’ contributions

CL drafted the manuscript, and developed the software. CL, IKG, ØB and TGL contributed to the conception and design of the study, and to the interpretation of results. All authors have read and approved the final version of the manuscript.

## Availability of data and materials

The TCGA BRCA data set analysed in this study is publicly available [15], and we downloaded both the BRCA RNA-Seq FPKM-UQ and the RPPA data set from the UCSC Xena Browser repository (http://xena.ucsc.edu) [24].

The Oslo 2 mRNA expression data are available from the GEO repository with accession number GSE58212 (https://www.ncbi.nlm.nih.gov/geo) [28]. The RPPA data from the Oslo 2 data set can be downloaded from Additional file 4 of the e-print [16].

The tailored graphical lasso has been implemented in the R package tailoredGlasso (https://github.com/Camiling/tailoredGlasso). R code for the simulations and data analyses in this paper is available at https://github.com/Camiling/tailoredGlassoAnalysis.

## Ethics approval and consent to participate

Not applicable.

## Consent for publication

Not applicable.

## Additional files

**Additional file 1** — Comparison of histograms of the non-zero prior partial correlation weights for the simulated data and the real multiomic data.

The histograms show how the distribution of the prior weights in our simulations resemble the distribution of the prior weights used in the multiomic applications. The histograms show the non-zero prior weights for the simulated data with partial correlations equal to (a) 0.02 and (b) 0.01, and the real genomic data from (c) the TCGA data set and (d) the Oslo 2 data set.

**Additional file 2** — The tailored graphical lasso graph for the TCGA BRCA RPPA data.

The graph found in the analysis of the TCGA BRCA data we did in this paper, using the RNA-seq data as prior information.

**Additional file 3** — The tailored graphical lasso graph for the Oslo 2 RPPA data.

The graph found in the analysis of the Oslo 2 data we did in this paper, using the mRNA data as prior information.

**Additional file 4** — List of genes with interactions in the TCGA BRCA tailored graphical lasso graph.

Each row contains the name of two genes whose nodes in the tailored graphical lasso graph have an edge.

**Additional file 5** — List of genes with interactions in the Oslo 2 tailored graphical lasso graph.

Each row contains the name of two genes whose nodes in the tailored graphical lasso graph have an edge.

**Additional file 6** — List of the gene interactions in the Oslo 2 tailored graphical lasso graph that the weighted graphical lasso was not able to find. Each row contains the name of two genes whose nodes have an edge in the tailored graphical lasso graph, but not in the weighted graphical lasso graph.

**Additional file 7** — List of the gene interactions in the TCGA BRCA tailored graphical lasso graph that the weighted graphical lasso was not able to find.

Each row contains the name of two genes whose nodes have an edge in the tailored graphical lasso graph, but not in the weighted graphical lasso graph.

**Additional file 8** — Comparison of evidence for edges found by the tailored graphical lasso and weighted graphical lasso for the Oslo 2 and TCGA data sets.

The number of edges found by the tailored graphical lasso and the weighted graphical lasso with evidence score ≥ 0.4 in different types of evidence. Note that one edge can have evidence of several types. See [23] for details on the evidence types and how the scores are calculated.

## References

1. Someren, E.v., Wessels, L., Backer, E., Reinders, M.: Genetic network modeling. Pharmacogenomics 3(4), 507–525 (2002)

2. Wang, Z., Gerstein, M., Snyder, M.: RNA-seq: a revolutionary tool for transcriptomics. Nature Reviews Genetics 10, 57–63 (2009)

3. Bergersen, L.C., Glad, I.K., Lyng, H.: Weighted lasso with data integration. Statistical Applications in Genetics and Molecular Biology 10(2011)

4. Lien, T.G., Borgan, Ø., Reppe, S., Gautvik, K., Glad, I.K.: Integrated analysis of DNA-methylation and gene expression using high-dimensional penalized regression: a cohort study on bone mineral density in postmenopausal women. BMC Medical Genomics 11, 24 (2018). doi:10.1186/s12920-018-0341-2

5. Kolaczyk, E.D.: Statistical Analysis of Network Data. Springer, New York, NY (2009)

6. Friedman, J., Hastie, T., Tibshirani, R.: Sparse inverse covariance estimation with the graphical lasso. Biostatistics 9, 432–441 (2008). doi:10.1093/biostatistics/kxm045

7. Banerjee, O., Ghaoui, L.E., d’Aspremont, A.: Model selection through sparse maximum likelihood estimation for multivariate Gaussian or binary data. Journal of Machine Learning Research 9, 485–516 (2008)

8. Meinshausen, N., Bühlmann, P.: High-dimensional graphs and variable selection with the lasso. The Annals of Statistics 34, 1436–1462 (2006)

9. Friedman, J., Hastie, T., Tibshirani, R.: Glasso: Graphical Lasso: Estimation of Gaussian Graphical Models. (2019). R package version 1.11. https://CRAN.R-project.org/package=glasso

10. Jiang, H., Fei, X., Liu, H., Roeder, K., Lafferty, J., Wasserman, L., Li, X., Zhao, T.: Huge: High-Dimensional Undirected Graph Estimation. (2020). R package version 1.3.4.1. https://CRAN.R-project.org/package=huge

11. Liu, H., Roeder, K., Wasserman, L.: Stability Approach to Regularization Selection (StARS) for high dimensional graphical models. In: Proceedings of the 23rd International Conference on Neural Information Processing Systems, vol. 2, pp. 1432–1440 (2010)

12. Foygel, R., Drton, M.: Extended Bayesian information criteria for Gaussian graphical models. In: Advances in Neural Information Processing Systems, vol. 23, pp. 604–612 (2010)

13. Li, Y., Jackson, S.A.: Gene Network Reconstruction by Integration of Prior Biological Knowledge. G3: Genes, Genomes, Genetics 5, 1075–1079 (2015). doi:10.1534/g3.115.018127

14. R Core Team: R: A Language and Environment for Statistical Computing. R Foundation for Statistical Computing, Vienna, Austria (2013). R Foundation for Statistical Computing. http://www.R-project.org/

15. Cancer Genome Atlas Network et al.: Comprehensive molecular portraits of human breast tumours. Nature 490, 61 (2012)

16. Aure, M.R., Jernström, S., Krohn, M., Vollan, H.K.M., Due, E.U., Rødland, E., Kåresen, R., Ram, P., Lu, Y., Mills, G.B., et al.: Integrated analysis reveals microRNA networks coordinately expressed with key proteins in breast cancer. Genome Medicine 7, 21 (2015). doi:10.1186/s13073-015-0135-5

17. Johansson, Å., Løset, M., Mundal, S.B., Johnson, M.P., Freed, K.A., Fenstad, M.H., Moses, E.K., Austgulen, R., Blangero, J.: Partial correlation network analyses to detect altered gene interactions in human disease: using preeclampsia as a model. Human Genetics 129(1), 25–34 (2011)

18. Schäfer, J., Strimmer, K.: An empirical Bayes approach to inferring large-scale gene association networks. Bioinformatics 21(6), 754–764 (2005)

19. Fujita, A., Sato, J.R., Garay-Malpartida, H.M., Yamaguchi, R., Miyano, S., Sogayar, M.C., Ferreira, C.E.: Modeling gene expression regulatory networks with the sparse vector autoregressive model. BMC Systems Biology 1(1), 1–11 (2007)

20. Szklarczyk, D., Gable, A.L., Lyon, D., Junge, A., Wyder, S., Huerta-Cepas, J., Simonovic, M., Doncheva, N.T., Morris, J.H., Bork, P., Jensen, L.J., Mering, C.v.: STRING v11: protein-protein association networks with increased coverage, supporting functional discovery in genome-wide experimental datasets. Nucleic Acids Research 47, 607–613 (2019)

21. Johansson, H.J., Socciarelli, F., Vacanti, N.M., Haugen, M.H., Zhu, Y., Siavelis, I., Fernandez-Woodbridge, A., Aure, M.R., Sennblad, B., Vesterlund, M., et al.: Breast cancer quantitative proteome and proteogenomic landscape. Nature Communications 10(1), 1–14 (2019)

22. Cancer Genome Atlas Network and others: Comprehensive molecular portraits of human breast tumours. Nature 490(7418), 61 (2012)

23. Von Mering, C., Jensen, L.J., Snel, B., Hooper, S.D., Krupp, M., Foglierini, M., Jouffre, N., Huynen, M.A., Bork, P.: String: known and predicted protein–protein associations, integrated and transferred across organisms. Nucleic Acids Research 33(suppl 1), 433–437 (2005)

24. Goldman, M.J., Craft, B., Hastie, M., Repečka, K., McDade, F., Kamath, A., Banerjee, A., Luo, Y., Rogers, D., Brooks, A.N., et al.: Visualizing and interpreting cancer genomics data via the Xena platform. Nature Biotechnology, 1–4 (2020). doi:10.1038/s41587-020-0546-8

25. Lee, H., Palm, J., Grimes, S.M., Ji, H.P.: The cancer genome atlas clinical explorer: a web and mobile interface for identifying clinical–genomic driver associations. Genome Medicine 7, 112 (2015)

26. Weizmann Institute of Science: GeneCards - The Human Gene Database. https://www.genecards.org Accessed 2019-04-11

27. Davis, S., Meltzer, P.: GEOquery: a bridge between the Gene Expression Omnibus (GEO) and BioConductor. Bioinformatics 14, 1846–1847 (2007)

28. Barrett, T., Troup, D.B., Wilhite, S.E., Ledoux, P., Rudnev, D., Evangelista, C., Kim, I.F., Soboleva, A., Tomashevsky, M., Marshall, K.A., et al.: NCBI GEO: archive for high-throughput functional genomic data. Nucleic acids research 37(suppl 1), 885–890 (2009)

