## Supplementary figures and images for "Tailored Graphical Lasso for Data Integration in Gene Network Reconstruction"

### Additional file 1

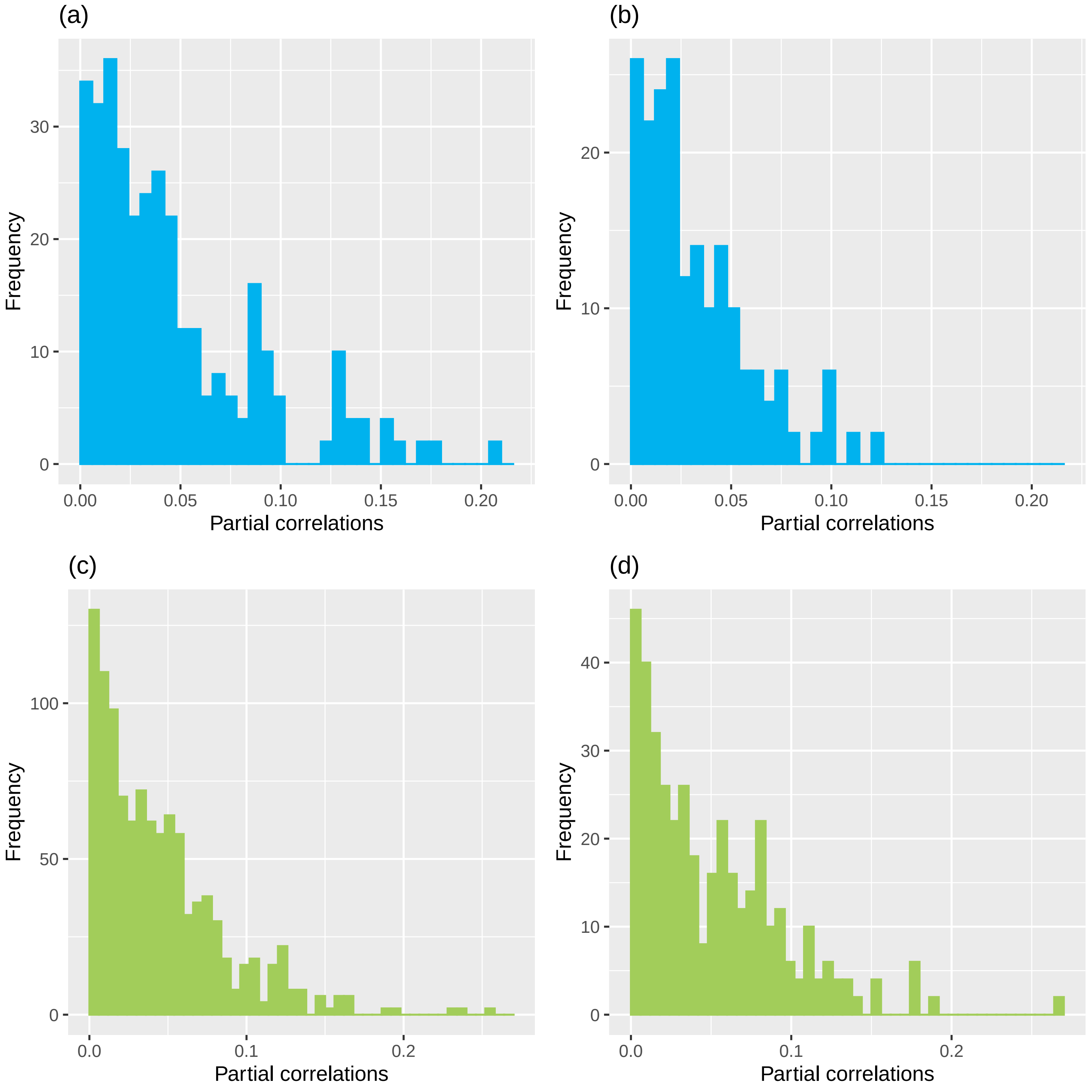

### Additional file 2

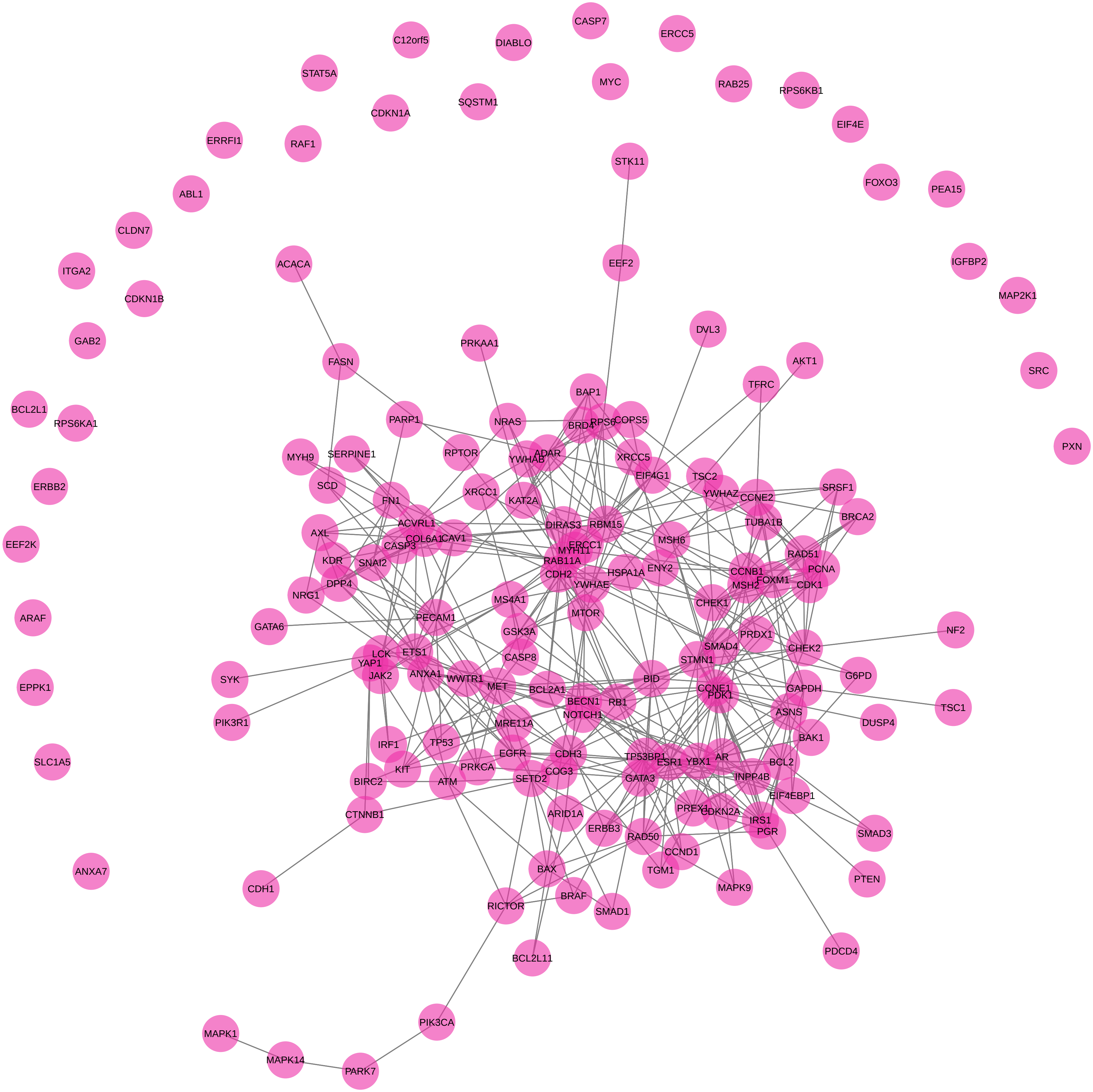

### Additional file 3

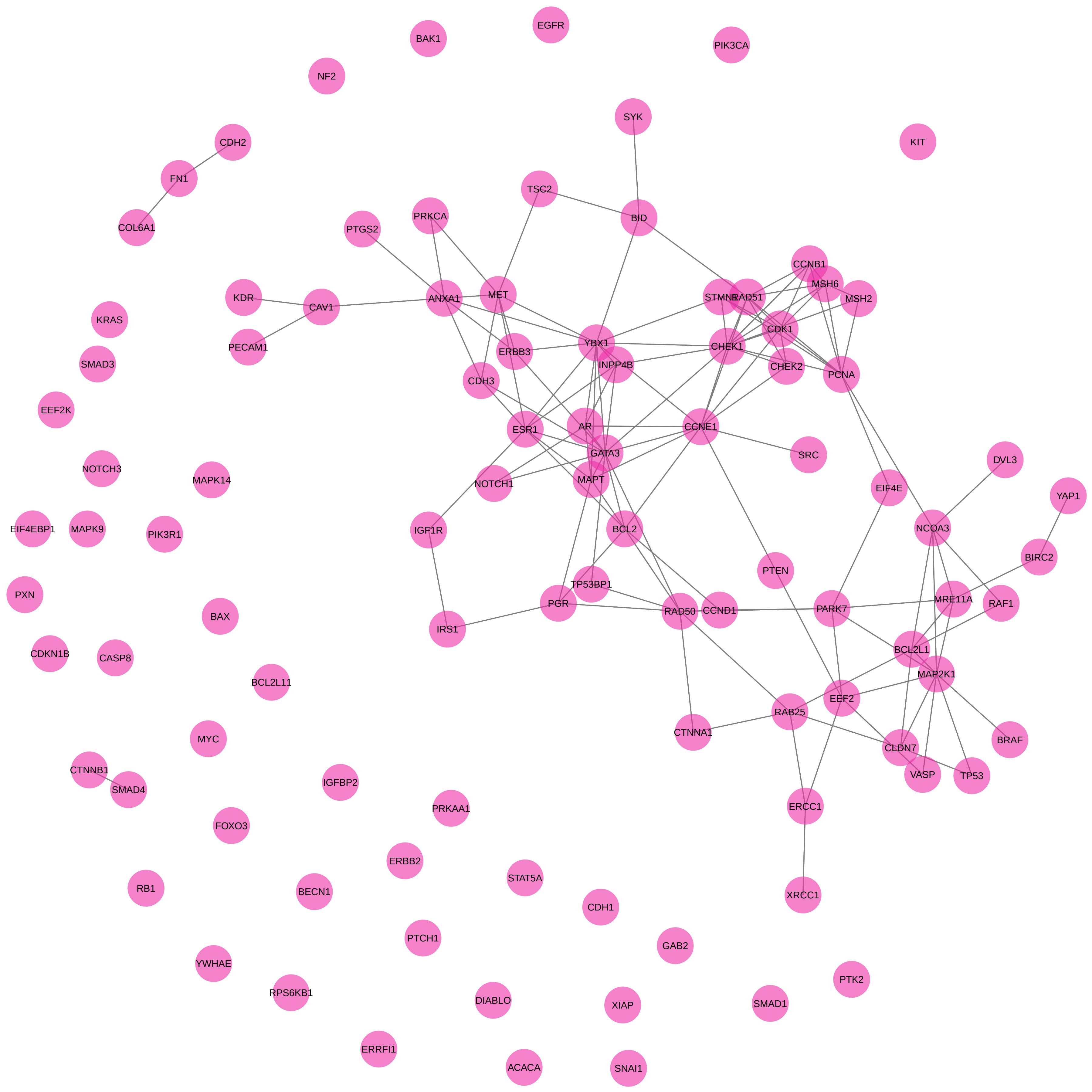
